# Predicting glaucoma prior to its onset using deep learning

**DOI:** 10.1101/828681

**Authors:** Anshul Thakur, Michael Goldbaum, Siamak Yousefi

**Author notes:** Corresponding Author: Siamak Yousefi, 930 Madison Ave., Suite 726, Memphis, TN 38163.

## Abstract

**Purpose:** To assess the accuracy of deep learning models to predict glaucoma development from fundus photographs several years prior to disease onset.

**Design:** A deep learning model for prediction of glaucomatous optic neuropathy or visual field abnormality from color fundus photographs.

**Participants:** We retrospectively included 66,721 fundus photographs from 3,272 eyes of 1,636 subjects to develop deep leaning models.

**Method:** Fundus photographs and visual fields were carefully examined by two independent readers from the optic disc and visual field reading centers of the ocular hypertension treatment study (OHTS). When an abnormality was detected by the readers, subject was recalled for re-testing to confirm the abnormality and further confirmation by an endpoint committee. Using OHTS data, deep learning models were trained and tested using 85% of the fundus photographs and further validated (re-tested) on the remaining (held-out) 15% of the fundus photographs.

**Main Outcome Measures:** Accuracy and area under the receiver-operating characteristic curve (AUC).

**Results:** The AUC of the deep learning model in predicting glaucoma development 4-7 years prior to disease onset was 0.77 (95% confidence interval 0.75, 0.79). The accuracy of the model in predicting glaucoma development about 1-3 years prior to disease onset was 0.88 (0.86, 0.91). The accuracy of the model in detecting glaucoma after onset was 0.95 (0.94, 0.96).

**Conclusions:** Deep learning models can predict glaucoma development prior to disease onset with reasonable accuracy. Eyes with visual field abnormality but not glaucomatous optic neuropathy had a higher tendency to be missed by deep learning algorithms.

## Introduction

While most of the application of deep learning models has been centered around glaucoma diagnosis for the screening purposes, the aim of our study is to evaluate the utility of deep learning models for prediction of glaucoma from color fundus photographs well prior to the manifestation of the clinical signs. We hypothesize that deep learning models, such as models that we propose, can uncover clinical and subclinical glaucoma-induced signs that may lead to improving our understanding of mechanisms underlying glaucoma.

Glaucoma is a heterogeneous group of disorders that represents the second leading cause of blindness overall, affecting up to 91 million individuals worldwide. ^1,2^ Glaucoma has multiple known risk factors including older age, African-American ethnicity, elevated intraocular pressure (IOP), and thinner central corneal thickness.^3,4^ However, subjects with one or more of these risk factors may or may not develop glaucoma making an accurate prediction challenging.^5^ Since glaucoma can be asymptomatic, its detection before significant vision loss is critical.^6^ Hence, methods for predicting glaucoma could have a significant impact on public health.

Dilated fundus photography provides convenient and inexpensive means for recording optic nerve head (ONH) structure and glaucomatous optic neuropathy (GON) assessment remains a gold standard for indicating the presence of glaucoma.^7,8^ However, manual assessment of the optic disc through fundus photographs for glaucoma screening requires significant clinical training, is highly subjective with currently limited agreement regarding results even among glaucoma specialists, and is labor intensive for application to the general population.^9,10^ Recent advances in artificial intelligence and deep learning models along with significant growth in available methods to record fundus photographs have shown promise and allowed the development of objective systems to assess the ONH through fundus photographs, thus leading to enhanced glaucoma diagnosis.^11-15^

Deep learning models require large clinically annotated training datasets to learn promising features from the images. Learning from data has advantages over predefined assumptions and rules to build the knowledge in machine learning classifiers. Several studies have shown that deep learning models can identify disease-induced signs to diagnose disease or identify the severity of disease from ophthalmic images with high accuracy in ocular conditions such as diabetic retinopathy, age-related macular degeneration, and glaucoma.^13,16-20^ Integrating deep learning models into portable fundus photography cameras or general practice may provide automated assessment of ocular conditions such as glaucoma and has a significant potential for providing affordable screening of at-risk populations and improving access to care.

## Method

### Subjects and Data

The fundus photographs of this study were obtained from the ocular hypertension treatment study (OHTS) after signing the data use agreement and receiving the institutional review board (IRB) approval. The study was conducted according to the tenets of Helsinki. The OHTS was a prospective, multi-center investigation (22 centers across the US) that sought to prevent or delay the onset of visual field loss in patients with elevated IOP (at moderate risk of developing glaucoma). All risk factors were measured at the baseline prior to disease onset and were collected for about 16 years (phases 1, 2). Hence, the longitudinal basis of the OHTS dataset allows development of models for predicting glaucoma prior to disease onset.

A total of 66,721 fundus photographs from 3,272 eyes of 1,636 subjects with normal appearing optic disc and normal visual field at the baseline visit were included. Ocular measurements and fundus photographs were collected every year over the course of the study. Details of the OHTS study and the procedure for identifying glaucoma are outlined in another study. ^21^ Figure 1 illustrates how fundus photographs were labeled and three datasets that we generated from a pool of 66,721 fundus photographs for developing three different deep learning models for detection and prediction of glaucoma.

**Figure 1.**
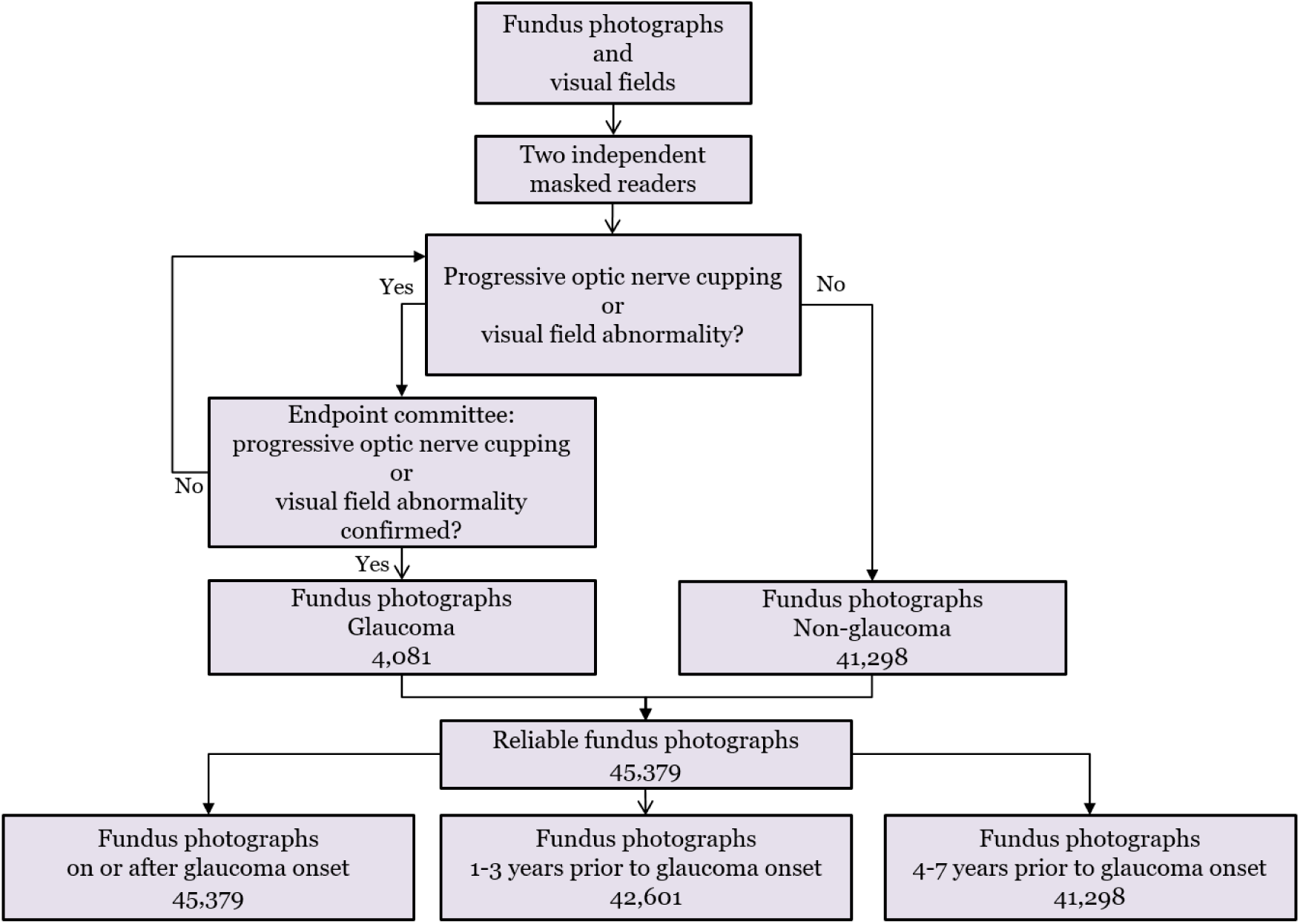
Flowchart of glaucoma identification and labeling. The eye is labeled as glaucoma based on either glaucomatous optic neuropathy (GON) or visual field abnormality. Reading center assignment should be further confirmation by an endpoint committee. Three datasets were selected from fundus photographs based on glaucoma onset date of each eye; one dataset for glaucoma diagnosis and two datasets for glaucoma prediction.

The first dataset included 45, 379 fundus photographs from non-glaucoma (throughout this manuscript, non-glaucoma refers to eyes with elevated IOP but normal structure and visual field, as defined in OHTS study) eyes and eyes with glaucoma. Out of 45,379 fundus photographs, 41,298 were from non-glaucoma eyes and the remaining 4,081 photographs were from eyes with glaucoma (determined based on GON or visual field abnormality). We called this the diagnosis dataset (Fig. 2, red arrow shows the time points of corresponding fundus photographs). From 4,081 fundus photographs from eyes with glaucoma, about 29% of these fundus photographs were labeled as glaucoma due to GON without any visual field abnormality, 22% of the photographs were labeled as glaucoma due to VF abnormality without any evidence of GON, and 49% of photographs were labeled as glaucoma due to existence of both GON and visual field abnormality.

**Figure 2.**
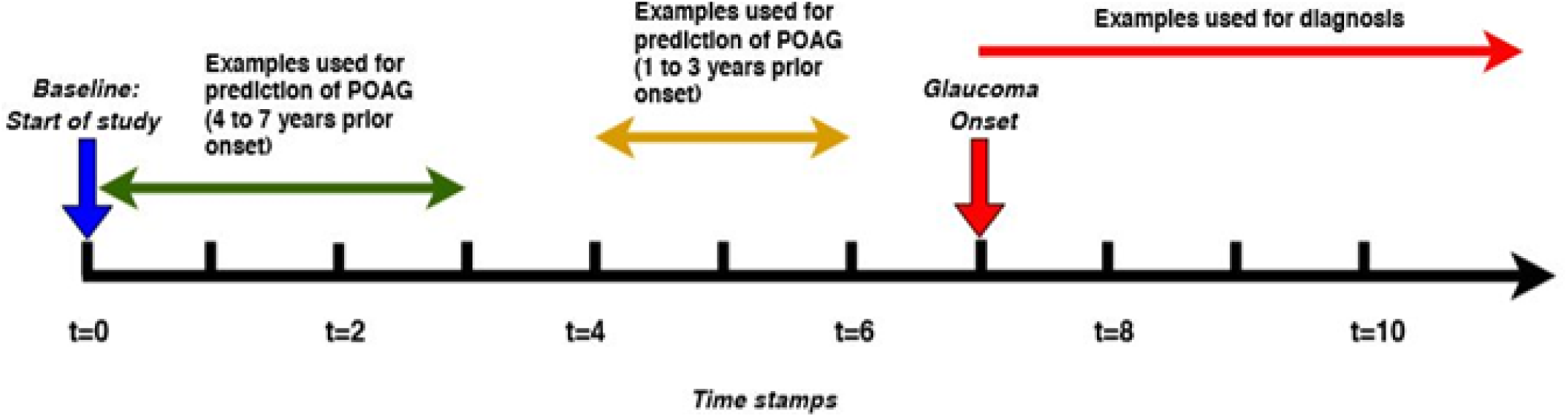
Eyes without any sings of glaucomatous optic neuropathy (GON) or visual field abnormality were followed for about 10 years and fundus photographs were collected annually. The onset time represents when a sample eyes was identified as glaucoma based on GON or visual field abnormality. Green arrow corresponds to fundus photographs collected 4-7 years prior to the date of glaucoma onset, yellow arrow represents fundus photographs that were collected 1-3 years prior to the date of glaucoma conversion, and red arrow corresponds to fundus photographs collected on or after the time that glaucoma onset was identified.

The second dataset included 42,601 fundus photographs from non-glaucoma eyes and eyes that eventually converted to glaucoma after about 1-3 years. We called this the “late prediction” dataset. The late prediction dataset includes 41,298 fundus photographs from non-glaucoma eyes and 1,303 fundus photographs from eyes that converted to glaucoma after 1-3 years (Fig. 2, yellow arrow shows the time points of corresponding fundus photographs). The third dataset included 42,498 fundus photographs from non-glaucoma eyes and eyes that eventually converted to glaucoma after about 4-7 years. We called this the “early prediction” dataset. Early prediction dataset included 41,298 fundus photographs from non-glaucoma eyes and 1,200 fundus photographs from eyes that eventually developed glaucoma after about 4-7 years (Fig. 2, green arrow shows the time points of corresponding fundus photographs). The same set of fundus photographs obtained from non-glaucoma eyes were used in all three datasets. We developed three different deep learning models using fundus photographs from these three datasets to assess the accuracy of models in predicting and detecting glaucoma.

#### Train, test, and validation datasets

We first selected 15% of the data for the final validating (re-testing). We then selected the rest of 85% of the data and used 5-fold cross-validation for training the models, hyperparameter selection, and testing. In each fold of the cross validation, the 80% of examples from each class were used for training while the remaining examples are used for testing. To avoid bias, all testing and validation were performed on the subject level rather than the eye level without any overlap between train, test, and validation.

### Image preprocessing

As mentioned before, OHTS fundus photographs were scanned from documented fundus photographs (printouts) and then saved in JPEG format. Therefore, fundus photographs from the OHTS dataset presented additional artifacts compared to common artifacts present in fundus photographs such as lighting conditions and effects of different environments and camera settings. Artifacts such as image deformation and presence of not-related labels on images is common in images from OHTS dataset. To mitigate some of the image quality issues, we performed contrast enhancement and applied Gaussian filtering to all fundus photographs. The photographs were cropped, normalized and resized to 224 × 224 × 3 (color format).

### Deep learning model

We used a computationally efficient convolutional neural network (CNN) architecture, referred to as MobileNetV2, ^22^ to develop our deep learning models. The trainable parameters in MobileNetV2 are about 1% of the parameters in competing models such as Inception-v3^23^ and ResNet-150^24^, making it a great choice for problems where computation resources and the training data are scarce.

The first deep learning model was trained on the “diagnosis dataset” in which fundus photographs had been taken on or after glaucoma onset and was used to classify a given fundus photograph as glaucomatous or non-glaucomatous (Fig. 1). One challenging aspect of this model is to identify fundus photographs that had been labeled as glaucoma due to visual field abnormality without any obvious sign of GON. This makes our diagnosis model more robust compared to previously developed models that only identify fundus photographs that have been labeled as glaucoma due to GON. Similarly, the other two models were trained for predicting glaucoma 1-3 and 4-7 years prior to disease onset. To the best of our knowledge, this is the first deep learning model that predicts glaucoma from fundus photographs several years prior to clinical functional or structural manifestation of the signs of glaucoma. Since the fundus photographs in the prediction models were collected from eyes prior to glaucoma onset and therefore without any obvious or clinical signs of glaucoma, prediction models are more challenging to develop than the diagnosis model.

### Training strategy

To train models, we used transfer learning. More specifically, MobileNetV2 was initialized with pre-trained weights that were initially obtained by training the model on ImageNet dataset. Transfer learning makes the convergence faster and provides a more effective classification performance when dealing with training data with small samples. We then fine tuned the general knowledge of image interpretation by learning from domain-specific OHTS fundus photographs.

As discussed previously, all three datasets had a greater number of fundus photographs from the non-glaucoma eyes compared to glaucoma eyes. In average, 9%, 3.1% and 2.9% of the fundus photographs in the diagnosis, first, and second prediction datasets were from glaucoma eyes, respectively. To address class imbalance issue, we performed data augmentation and applied balanced data sampling for batch creation. We performed random horizontal and vertical flips, rotations and randomly changing the hue, saturation and contrast of the training fundus photographs. After augmentation, during training, the same number of photographs from both classes were selected for each mini batch. A batch size of 64 images, cross-entropy loss function, and Adam optimizer with a fixed learning rate of 0.001 were used for training the models. To avoid overfitting, weight decay of 0.0001 is used on all layers of the model. All programs were implemented in Python with a backend of PyTorch.

### Deep learning interpretation

To identify the regions of the fundus photographs that drive the deep learning model to assign an image to glaucoma or non-glaucoma groups, we used gradient-weighted class activation maps. ^21^ The activation maps use the final convolutional layer of a CNN to produce a coarse localization map of the driving regions of the fundus photographs used for diagnosis.^21^ While activation maps can be used to validate deep learning models to verify clinically relevant regions are assess for diagnosis, they can be used to discover potentially novel biomarkers for the disease as well.

### Statistical Analyses

Models were tested using 5-fold cross-validation datasets (independent from training datasets) and validated (re-tested) using the held-out dataset. The performance of each model was assessed using AUC. The method of DeLong et al.^25^ was used to compare the AUC of different models. All statistical analyses were performed in Python.

## Results

About 24% of the fundus photographs in the OHTS dataset had extreme artifacts and were excluded from the study. Figure 3 shows the activation maps obtained from three glaucomatous and three normal fundus images using the model trained on the diagnosis dataset. As can be seen, activation maps confirm that optic cup and rim were the most important regions in the input fundus photographs of eyes without glaucoma (first row) and eyes with glaucoma (third row).

**Figure 3.**
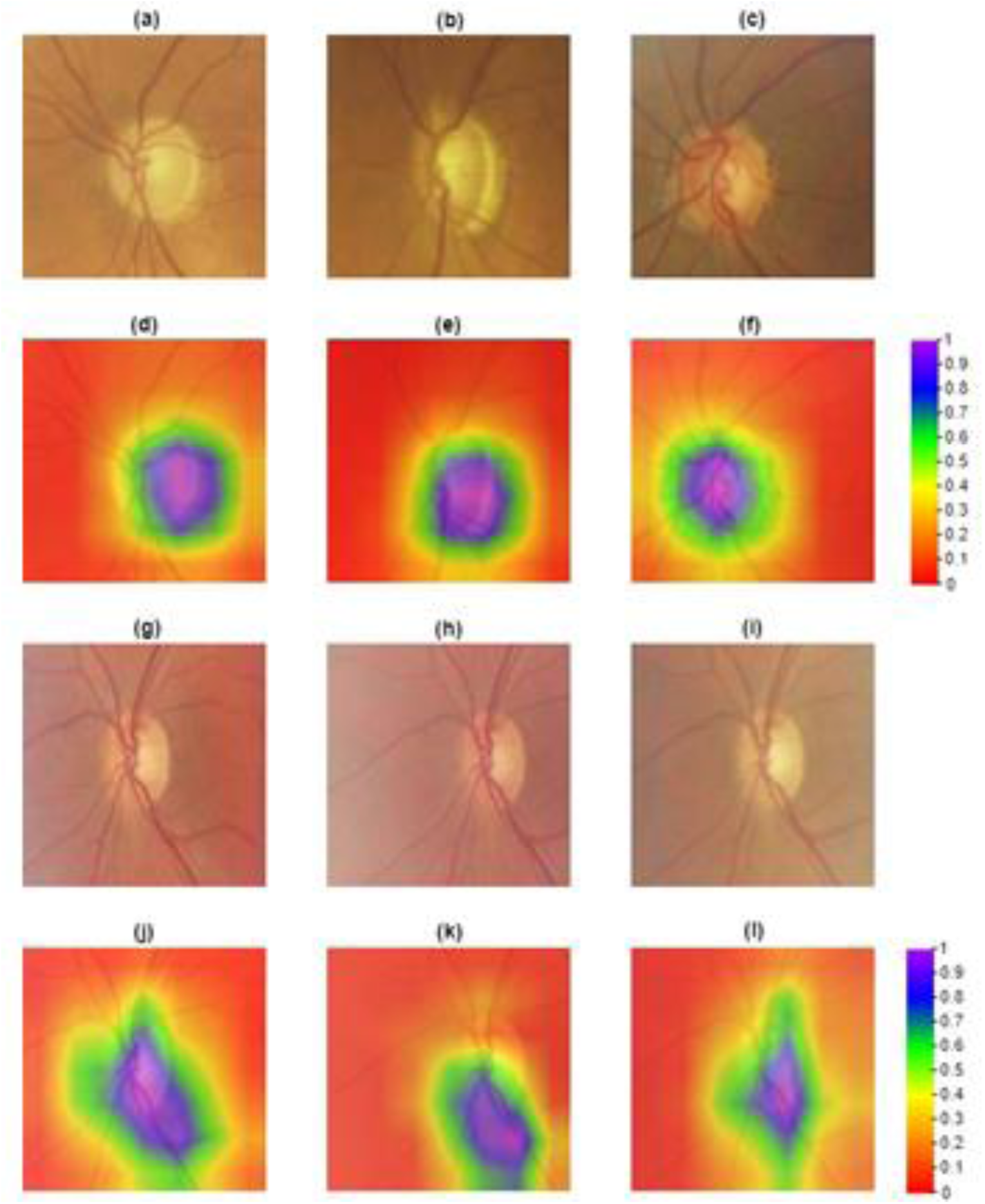
Activation maps representing regions that are most promising for the deep learning model to make a diagnosis. First row: fundus photographs from non-glaucoma eyes. Second row: activation maps of fundus photographs of non-glaucoma eyes (shown in the first row). Third row: fundus photographs from eyes with glaucoma. Fourth row: activation maps of fundus photographs of eyes with glaucoma (shown in the third row).

Figure 4 shows the ROC curves of the three deep learning models that were re-tested using the held-out subset of the diagnosis, first prediction, and second prediction models, respectively. The AUC of the deep learning model for making diagnosis was 0.945 (95% confidence interval; CI of 0.93 to 0.96). The AUC of the deep learning model on the first prediction dataset and second prediction dataset were 0.88 (0.86, 0.91) and 0.77 (0.75, 0.78), respectively.

**Figure 4.**
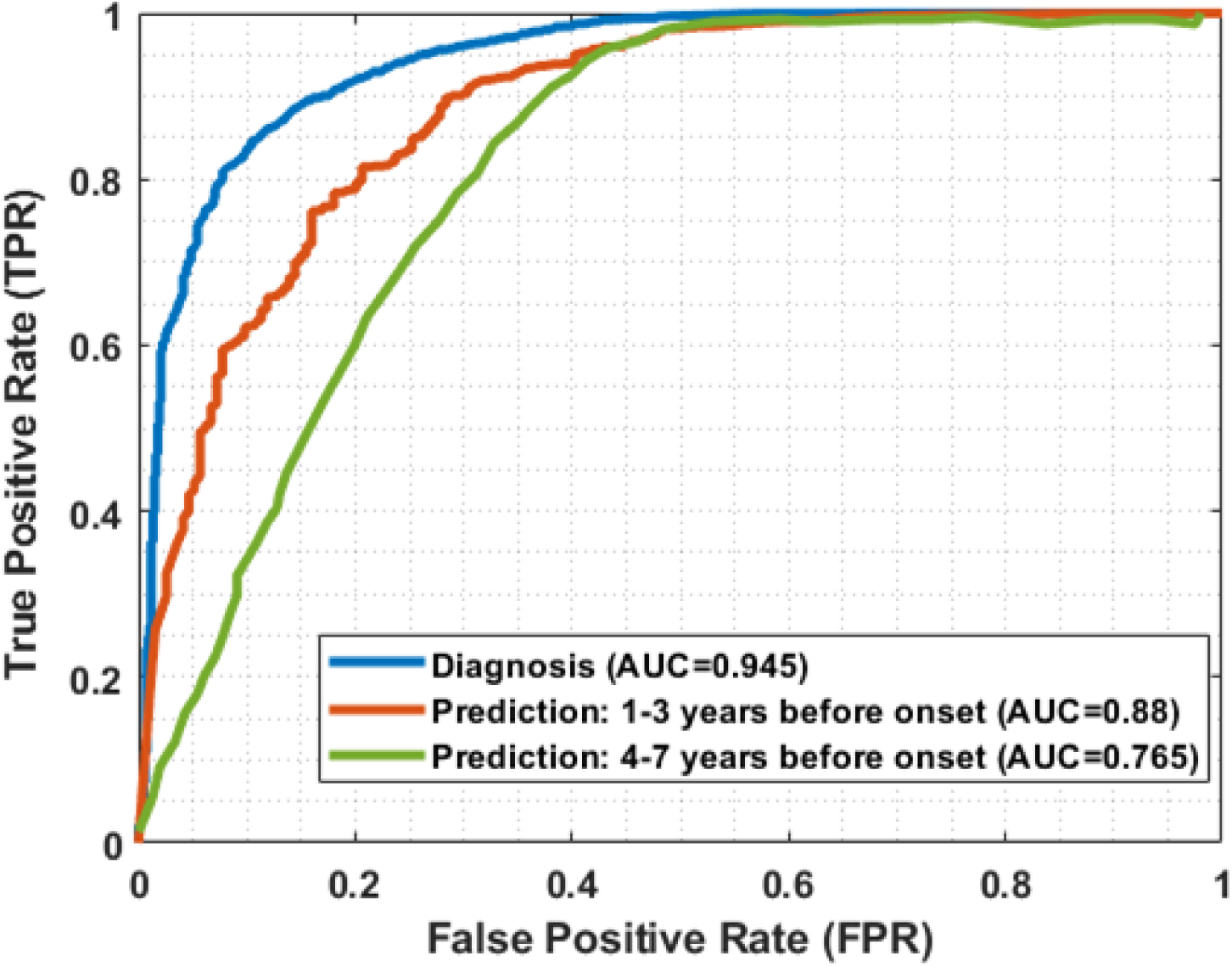
Receiver operating characteristic (ROC) curves of the prediction and diagnosis models. Green curved belongs to the model that predicts glaucoma 4-7 years prior to the onset of the disease, red curve predicts glaucoma 1-3 years prior to the onset of the disease, and blue curve shows diagnosing glaucoma on or after onset.

For the task of diagnosis, the AUC was improved to 0.97 (0.96, 0.98) on re-testing fundus photographs that were labeled as glaucoma due to GON. However, as expected, the AUC decreased to 0.88 (0.86, 0.89) when we tested the diagnostic deep learning model using the fundus photographs that were labeled as glaucomatous due to abnormal visual field without GON.

## Discussion

Unlike previous studies with the emphasis on diagnosis, we proposed models for prediction of glaucoma prior to disease onset. The proposed models showed consistent performance in predicting glaucoma development 1-3 years and 4-7 years prior to the disease onset. There are several studies that have proposed deep learning for identifying glaucoma from fundus photographs. However, all these methods are centered around glaucoma diagnosis from fundus images that have been collected several years after the initial onset of the disease while in this study, we tackled a more challenging task of predicting glaucoma prior to manifestation of clinical signs.

Fundus photography provides a simple, cheap and more portable means for screening in underserved populations by non-physicians thus improving access to care. Recently, several deep learning approaches have been proposed for detecting glaucoma from fundus photographs. Raghavendra et al.^11^ developed a deep learning model composed of 18 layers trained and tested on abount 1,500 fundus photographs and were able to reach an accuracy of 98% for diagnosing glauocma. The AUC of the six-layer deep learning model for glauocma diagnosis proposed by Xiangyu et al. ^12^ was 0.83 and 0.88 on two different datasets, respectively. Li et al. ^13^ applied a deep learning model using Inception-v3^23^ architecture on a large dataset with about 40,000 fundus photographs and achieved an AUC of about 0.99 in detecting referable glauocma defined based on GON. In another study, Christopher et al.^14^ developed several deep learning architectures and used a dataset with about 14,000 fundus photographs and achieved an AUC of 0.91 in identifying GON eyes. Nourizifard et al.^15^ developed multiple deep learning models and used relatively small datasets with total of about 500 fundus photographs and achieved an accuracy of 92%. The reported accuracy of the previous deep learning models to diagnose glaucoma ranges from 0.83 to 0.98.

Except studies that were conducted by Li et al. ^13^ and Christopher et al.^14^, the rest of the studies have used relatively small datasets of fundus photographs. Therefore, it is challenging to generalize their conclusions. However, we have used 66,721 fudnus photographs for training and tesing models to develop robus models and generate reproducible outcome. Aside from large datasets that are required to successfully develop deep learning models, validation is an important step. We have used three valiations steps; cross-validation, held-out subset, and visualization of promising features through activation maps. Thererfore, our models are robust and likely will be generalized to new data.

While the diagnostic AUC of our model was 0.94, the diagnostic accuracy of the model proposed by Raghavendra et al. was 0.98 and the diagnostic AUC of the model proposed by Li et al. was 0.99. The lower AUC of our model could be due to several reasons. First, fundus photographs of the OHTS study have lower quality compared to the fundus photographs used in other studies. Second, about 22% of the fundus photographs in the OHTS study were labeled glaucoma due to visual field abnormity without apparent GON. Therefore, it is more challenging to identify glaucomatous eyes without GON from fundus photographs. In fact, our supplemental analysis showed that the AUC will increase to 0.97 if we use only fundus photographs that were labeled as glaucoma due to GON.

Our study used a fully automated model to identify promising glaucoma-induced features (signs) while several previous studies used semi-objective hand-engineered features^26,27^, and therefore adopted ad hoc rules. In fact, we showed that using fully automated CNNs, we can achieve an AUC of 0.88 for predicting glaucoma 1-3 years prior to the manifestations of the clinical signs. Although the human expert may not identify subclinical changes in the optic nerve, the AUC of 0.88 highlights that the optic nerve has gone through subtle changes prior to the clinical manifestation of the disease. These subtle changes may be identified by a deep learning model but not by a human expert.

The OHTS dataset that was used in our study had several strengths; participants were recruited from 22 centers across the US, thereby reducing the idiosyncrasies of a local databases; reading centers had access to well-trained certified readers, data was collected, annotated and cured very well, disease onset was confirmed by repeating image and data collection, and endpoint committees further confirmed disease onset based on guidelines. Nevertheless, it had several limitations too. One limitation was poor quality of the scanned photographs from documented optic nerve printouts. However, even in the presence of poor quality and high variability, the diagnostic AUC was 0.94, which is promising. The other limitation was the smaller number of eyes with glaucoma compared to non-glaucoma eyes. However, this is a common problem in many healthcare domain and not specific to this study. To address this issue, we used a deep learning model with relatively small number of parameters, performed data augmentation and conducted data sampling for balanced batch creation. Another limitation is that OHTS dataset collected from a restricted clinical trial and eyes with elevated IOP therefore do not represent general population and real settings. Future studies with independent datasets from eye clinics may further verify the proposed models.

Despite the limitations, our study showed that deep learning models were sufficiently sensitive to predict eyes that will convert to glaucoma from baseline images. This study is an example of how the high performance of current deep learning classifiers allows us to imagine other important predictions that human observers are unable to make from routinely obtained medical images. These methods may open new eras in developing deep learning models for predicting glaucoma more accurately as well as other blinding eye diseases well in advance of the onset of the disease in order to identify at-risk population. Out study may also identify previously unknown signatures of the disease development.

## Acknowledgment

We thank Dianna Johnson for her assistance. This work was supported by NIH R21EY030142 (SY) and in part by an unrestricted grant from Research to Prevent Blindness (RPB), New York, (SY). The funders had no role in study design, data collection and analysis, decision to publish, or preparation of the manuscript.

